# MetaKnogic-Alpha: A Hyper-Relational Knowledge Base for Grounded Metabolic Reasoning

**DOI:** 10.64898/2026.02.05.704050

**Authors:** Pengtao Dang, Paveethran Swaminathan, Tingbo Guo, Changlin Wan, Sha Cao, Chi Zhang

## Abstract

The exponential trajectory of biomedical literature has precipitated a fundamental “synthesis gap” in metabolic research, where critical mechanistic insights remain fragmented across hundreds of thousands of disjointed full-text articles, preventing the consolidation of a global mechanistic view. Here, we present **MetaKnogic-Alpha**, a foundational mechanistic knowledge substrate designed to bridge this gap by transforming unstructured literature into a navigable, logic-based resource. MetaKnogic-Alpha synthesizes over 100K full-text articles into a hyper-relational hypergraph structure, preserving the n-ary relational logic inherent in complex metabolic pathways. To ensure biological rigor, we implemented a hierarchical discovery protocol: an autonomous reasoning agent first enriches query nomenclature for domain-specific precision, followed by a multi-hop topological expansion within the hypergraph to surface functional neighbors, such as enzymatic co-factors and distal regulators, often lost in traditional search paradigms. Crucially, the system subjects all literature-derived insights to a deterministic biochemical grounding against a curated metabolic reaction network, significantly mitigating the risk of probabilistic hallucinations common in standalone generative models. In rigorous benchmarking, MetaKnogic-Alpha achieved a mechanistic accuracy of 0.98 in scenarios where supporting evidence was present, providing a robustly attributable audit trail back to the primary literature via PubMed Central Identifiers. We designate this primary release as “alpha” to establish the foundational architectural logic for a burgeoning million-scale resource. By compressing the synthesis of thousands of papers from a multi-month manual effort into several hours of automated discovery, MetaKnogic-Alpha serves as a high-fidelity research companion that augments the human expert’s ability to resolve complex metabolic interactions and identify novel therapeutic drivers in precision oncology.

## 1 Introduction

The architecture of cellular metabolism is characterized by its profound non-linearity and systemic interconnectedness[1, 2]. Every enzymatic transformation, metabolite flux, and regulatory signaling node exists within a dense web of stoichiometric and thermodynamic constraints that span across organelles, cell types, and physiological states[3, 4, 5, 6, 7, 8]. In the current era of high-throughput multi-omics and precision oncology, the bottleneck to transformative discovery is no longer a lack of data, but rather a “synthesis gap”. With the exponential growth of the biomedical literature which now exceeding several thousand publications annually in the field of metabolism alone, the ability of the human researcher to consolidate fragmented experimental findings into a cohesive mechanistic narrative has been outpaced[9, 10, 11, 12].

Critical insights regarding a specific metabolic vulnerability in cancer or a novel regulatory feedback loop are often geographically dispersed across disjointed full-text articles. Consequently, the field faces a “fragmentation of truth”, where the global mechanistic context required for precision medicine remains latent and inaccessible within unstructured text[13, 14, 15]. While Large Language Models (LLMs) have emerged as powerful tools for textual synthesis, their utility in the rigorous domain of metabolism is severely limited by a lack of biological grounding and a propensity for “probabilistic hallucinations”. Standalone generative agents, despite their linguistic fluency, lack an innate understanding of biochemical stoichiometry and are unable to provide a verifiable audit trail for their assertions[16, 17, 18, 19].

To bridge this gap, the field has recently explored Retrieval-Augmented Generation (RAG) frameworks[20]. However, standard RAG implementations frequently suffer from “contextual fragmentation”, as they typically retrieve isolated text chunks based on superficial lexical similarity[21, 22]. In the context of metabolism, where a single reaction involves multiple *n*-ary participants and distal regulatory inputs, flat vector-based retrieval fails to capture the topological neighborhood of a biological event[23, 24, 25].

Recent advancements in Graph-augmented RAG (Graph-RAG)[26] and Hypergraph-based retrieval frameworks (Hypergraph-RAG) [27] have sought to mitigate this fragmentation by modeling knowledge as complex relational networks. However, these existing methodologies often encounter a “computational ceiling” when applied to million-scale datasets. Current Graph-RAG pipelines are frequently characterized by prohibitive operational costs and extreme latency, as the iterative graph-building, graph-traversal and global indexing processes can consume weeks or months of processing time for even moderate corpora[28, 29, 30, 31, 32]. This temporal and financial overhead renders them impractical for real-time discovery or for the dynamic ingestion of the ever-expanding biomedical literature.

Furthermore, many of these frameworks function as passive retrieval pipelines rather than integrated knowledge resources; they often lack the deterministic biochemical constraints and agentic reasoning required to ensure the stoichiometric integrity of a discovered pathway[33, 34, 35]. There is an urgent need for a high-fidelity knowledge substrate that can not only preserve the structural logic of metabolism but also provide a computationally efficient and verifiable interface for navigating the vast scale of the literature.

Here, we present **MetaKnogic-Alpha**, see Fig. 1, a foundational mechanistic knowledge substrate designed for automated metabolic discovery and high-resolution literature synthesis. MetaKnogic-Alpha represents a paradigm shift from passive, high-latency retrieval to an active, high-efficiency discovery infrastructure. By utilizing an asynchronous, distributed ingestion engine, we achieve a three-order-of-magnitude reduction in processing time, effectively compressing a multi-month synthesis timeline into several hours. We designate this primary release as “alpha” to establish the fundamental architectural logic of a burgeoning million-scale resource. By transforming over 100K full-text articles into a navigable, hyper-relational knowledge structure and grounding all findings within a deterministic metabolic reaction network, MetaKnogic-Alpha provides a “global mechanistic view” that was previously unattainable for the human expert without sacrificing computational feasibility.

**Figure 1.**
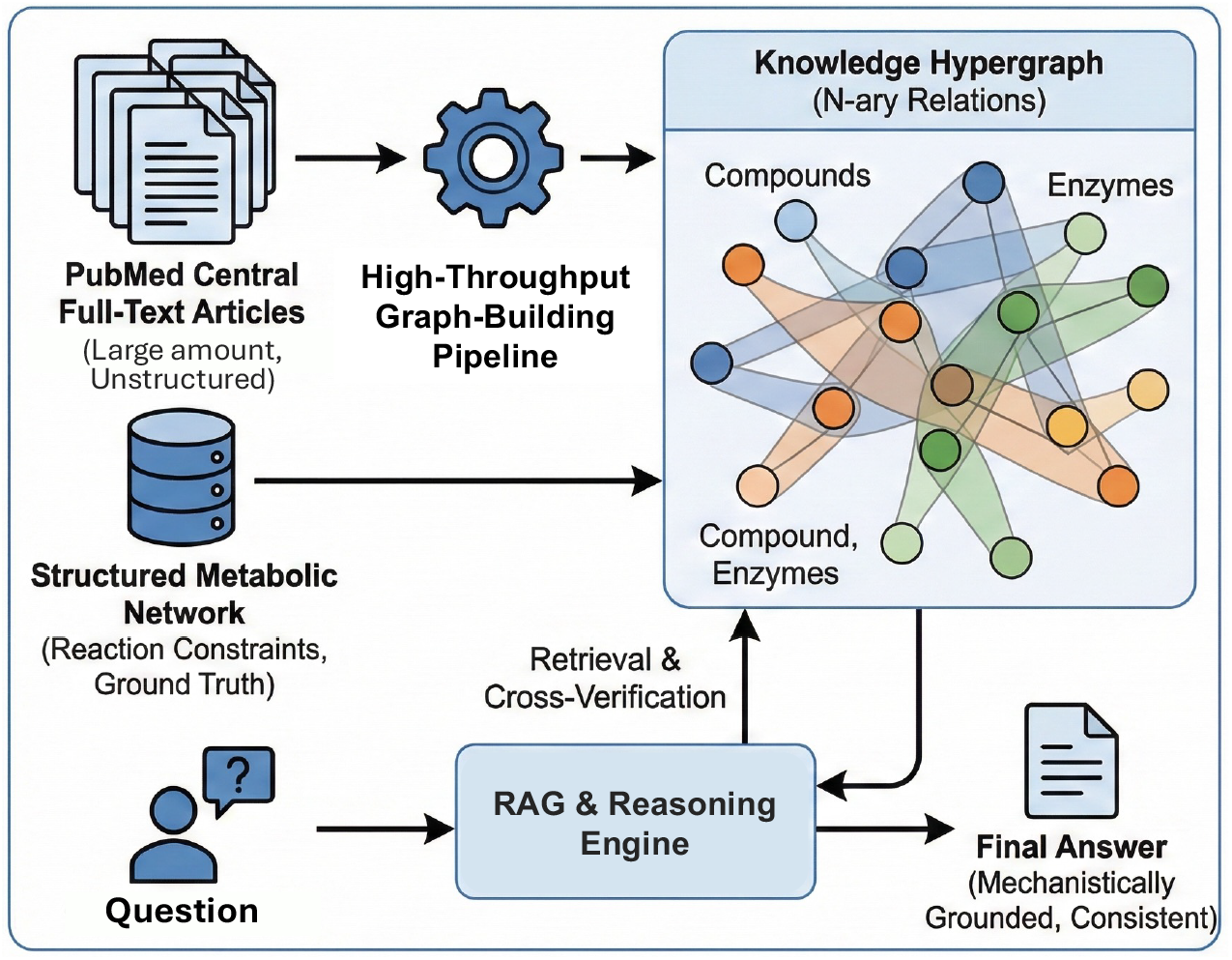
Overview of the MetaKnogic-alpha Discovery Framework. The MetaKnogic-alpha architecture integrates unstructured literature with deterministic biochemical logic to facilitate automated metabolic discovery. Large-scale, unstructured full-text articles are processed through a high-throughput pipeline to instantiate a Knowledge Hypergraph, capturing complex *n*−*ary* relations. To ensure biological rigor and mitigate probabilistic hallucinations, the framework incorporates a Structured Metabolic Network as a deterministic ground truth. When a mechanistic query is issued, the RAG & Reasoning Engine performs a multi-hop retrieval and cross-verification against the hypergraph and network constraints, yielding a Final Answer that is robustly attributable and mechanistically grounded in established biochemical principles.

The **MetaKnogic-Alpha** framework is built upon three core pillars of innovation:

1. **Agentic Query Enrichment and Semantic Discovery:** Utilizing a specialized reasoning agent, the system refines and expands abstract user inquiries into precise biomedical nomenclature. These queries are projected into a 768-dimensional latent space, identifying dense semantic “seeds” that ensure the discovery process is anchored in contextual meaning rather than lexical coincidence.
2. **Multi-Hop Topological Expansion:** Unlike “flat” retrieval systems, MetaKnogic-Alpha utilizes a literature hypergraph to execute multi-hop breadth-first traversals. This allows the system to surface first- and second-order functional neighbors, that are functionally coupled in vivo but may be disjointed across the literature corpus.
3. **Deterministic Metabolic Grounding:** To ensure biochemical plausibility, all literature-extracted insights are subjected to structural validation against a deterministic metabolic network. This “biochemical sanity check” filters out stochastic noise and ensures that the system’s reasoning adheres to established metabolic laws.

MetaKnogic-Alpha is not intended to replace the intuition of the human scientist; rather, it serves as a high-fidelity research companion that organizes the collective intelligence of the biomedical community into a coherent, logical, and attributable substrate for discovery.

## 2 Results

### 2.1 Scalable Construction of a Domain-Agnostic Knowledge Hypergraph

The transition from probabilistic text generation to structured mechanistic reasoning requires a fundamental re-building of how information is ingested and organized. Traditional RAG frameworks are frequently constrained by synchronous, stream-based processing models, which impose severe limitations on document throughput and operational scalability[31, 27]. To address these systemic bottlenecks, we developed **MetaKnogic-Alpha**, featuring a decoupled, high-throughput batch processing architecture designed for the mass-scale transformation of unstructured corpora into high-order relational structures.

Our pipeline introduces a strategic paradigm shift by replacing real-time inference with a parallelized asynchronous orchestration layer. This architecture executes a multi-stage refinement protocol: structural decomposition of dense documents, high-fidelity entity disambiguation, and the automated instantiation of *n*-ary hyperedges. As shown in Table 1 By leveraging an asynchronous execution strategy, the system circumvents the latency overhead inherent in sequential API calls. This architectural optimization resulted in a 50% reduction in total operational costs while enabling the processing of over 100K full-text articles, a corpus size previously deemed cost, prohibitive for academic research environments. Whereas conventional methods would require several months to process and ingest such a volume of data[27, 28], the MetaKnogic-Alpha pipeline successfully compressed this temporal requirement into hours, demonstrating a several-hundredfold increase in computational efficiency.

**Table 1:** System Performance Metrics and Hypergraph Structural Statistics.

| Architectural Feature | Observed Performance / Metric |
| --- | --- |
| Dataset Scale (Full-Text) | 100k+ Articles (Scalable to $10^6+$) |
| Ingestion Workflow | Optimized High-Throughput Offline Batching |
| Temporal Efficiency | Weeks vs. Months (Baseline) |
| API Expenditure Optimization | 50% Cost Reduction |
| System Utility | Integrated Web-UI & Dynamic Hypergraph Visualization |
| Knowledge Density | $N$-ary Multi-entity Relationship Extraction |

Crucially, the MetaKnogic-Alpha infrastructure is not merely a backend processing engine but a comprehensive system designed for scientific auditability. We integrated a high-performance web-based interface and an interactive visualization suite as fundamental components of the system’s utility. These tools move beyond simple data presentation, offering a transparent “audit trail” for every retrieved answer. By visualizing the underlying hypergraph connectivity and the specific literature-derived evidence paths, researchers can verify the mechanistic logic of the AI’s responses. This interpretability is essential for professional applications in scientific and research contexts, where the provenance of information is as critical as the information itself.

The resulting hypergraph structure currently encompasses millions of unique entities and hyperedges, capturing the complex, multi-entity relationships that define modern scientific consensus. The system is architected for seamless horizontal scaling; we demonstrate that the current 100K-document foundation can be readily expanded to million-scale coverage without a corresponding exponential increase in computational overhead. This scalability ensures that MetaKnogic-Alpha serves as a robust infrastructure for any knowledge-intensive domain, facilitating the synthesis of massive, fragmented data into a cohesive and interactive knowledge landscape.

### 2.2 KnowledgeHyperGraph-RAG Outperforms GPT-5.1 in Mechanistic Question Answering

The clinical utility of biomedical AI is fundamentally limited by the “probabilistic bottleneck”, where general-purpose models prioritize linguistic fluency over mechanistic accuracy[24, 33]. To quantify the impact of our hyper-relational architecture, we benchmarked MetaKnogic-Alpha against GPT-5.1 using a subset of the PubMedQA dataset[36]. From an initial pool of approximately 210,000 question-answer pairs, we identified approximately 1,000 high-complexity queries with direct PMCID overlap in our 100k-article full-text knowledge base. This selection criteria ensured that the evaluation focused on the system’s ability to synthesize specific, evidence-backed mechanistic insights rather than relying on the general training data of the foundation model.

As shown in Table 2The performance gap between the standalone LLM and the MetaKnogic-Alpha framework is substantial. For the entire evaluation set, GPT-5.1 (LLM-only) achieved an accuracy of 0.63 with a recall of 0.36. In contrast, the MetaKnogic-Alpha system reached an accuracy of 0.96 and a recall of 0.61, representing a significant leap in both precision and information retrieval. The advantage of hypergraph-grounding becomes even more pronounced when analyzing “paper hit questions”, queries for which specific supporting documents were identified within the hypergraph. In these scenarios, MetaKnogic-Alpha attained a near perfect accuracy of 0.98, compared to 0.64 for the standalone LLM. With a paper hit rate of 78%, the system demonstrates a robust capability to bridge the gap between unstructured query intent and verifiable literature evidence.

**Table 2:** Comparative Benchmarking on PubMedQA: GPT-5.1 vs. MetaKnogic-Alpha.

| Metric | GPT-5.1 | MetaKnogic-Alpha | $\Delta$ Improvement |
| --- | --- | --- | --- |
| *All Questions ($N \approx 1,000$)* |  |  |  |
| Accuracy | 0.63 | **0.96** | +52.4% |
| F1-Score | 0.38 | **0.43** | +13.2% |
| Recall | 0.36 | **0.61** | +69.4% |
| *Paper Hit Questions (78% hit rate)* |  |  |  |
| Accuracy | 0.64 | **0.98** | +53.1% |
| F1-Score | 0.40 | **0.52** | +30.0% |
| Recall | 0.39 | **0.62** | +59.0% |
| *Not Paper Hit Questions (22% of total)* |  |  |  |
| Accuracy | 0.56 | **0.81** | +44.6% |
| F1-Score | 0.24 | **0.30** | +25.0% |
| Recall | 0.18 | **0.27** | +50.0% |

A critical innovation of MetaKnogic-Alpha is its resilience against biological hallucinations. While GPT-5.1 is prone to generating biochemically implausible assertions when queried on difficult cases, our system enforces “mechanistic grounding” by requiring every generated response to be supported by a traversable *n*-ary evidence path. This addresses the identified barrier where medical LLMs fail to reliably surface deep mechanistic relations. By visualizing the underlying hypergraph connectivity, including the specific entities and relationships extracted from full-text literature, MetaKnogic-Alpha provides a transparent “audit trail” that allows clinicians to verify the logic of the AI’s conclusions. This interpretability is vital for tumor board applications, where the provenance of a therapeutic recommendation is as critical as the recommendation itself.

The superiority of the MetaKnogic-Alpha framework is largely attributed to the ingestion of full-text articles rather than mere abstracts. Full-text documents contain the granular experimental results and secondary metabolic effects required to resolve multi-step reasoning tasks that are absent from summary-level data. By representing these interactions as hyperedges, the system maintains the integrity of multi-entity relationships, such as complex enzyme-metabolite-pathway assemblies, which are often lost in traditional binary graph representations. This structural fidelity ensures that the system remains logically consistent even when scaling to million-document repositories, providing a scalable solution for precision medicine that significantly exceeds the capabilities of current state-of-the-art foundation models.

### 2.3 Integration of Structured Metabolic Networks for Logical Grounding

Literature-derived hypergraphs capture breadth but are ultimately observational. To enforce mechanistic consistency, we integrated a curated metabolic reaction network into the MetaKnogic-Alpha architecture as a biochemical grounding layer[3, 4]. This dual-pathway design constrains reasoning to literature evidence that is compatible with stoichiometric structure and curated reaction directionality.

Upon receiving a query, the system performs LLM-driven query expansion to map user language to metabolic ontology terms, then traverses the internal metabolic network (reactions, compounds, enzymes) to extract a relevant sub-network (e.g., directionality, substrate–product roles, and neighborhood context). The resulting biochemical context is injected into the prompt and used to steer hypergraph retrieval toward mechanistically plausible evidence rather than keyword proximity.

#### Effect of metabolic grounding

Metabolic grounding is most evident in our “Mechanistic Error Correction” case studies. Without biochemical constraints, standalone baselines sometimes propose reaction sequences that violate curated directionality or imply unsupported gene– reaction and substrate/product relationships. Grounding reduces these unsupported assertions by restricting retrieval and synthesis to network-supported relations under directionality and compartment constraints. It also improves multi-hop coherence by prioritizing mechanistically connected neighborhoods (shared metabolites/reactions and pathway-consistent chains) and increases traceability by anchoring outputs to the seed entities and traversed edge types. In scenarios involving enzyme inhibition and secondary metabolite accumulation, the grounded system identified directionality-constrained dead ends and infeasible bypasses that were overlooked by LLM-only models, preventing incorrect mechanistic narratives.

This integration very evidently demonstrated that the value of the 100k-article full-text hypergraph is significantly amplified when coupled with structural expert knowledge. By enforcing a “biochemical sanity check” on every retrieval, MetaKnogic-Alpha moves beyond the limitations of standard RAG systems, providing a robust and scalable infrastructure for mechanistic discovery that is as rigorous as it is broad.

### 2.4 Interactive Visualization and Mechanistic Interpretability

The clinical adoption of large language models is frequently hindered by a lack of transparency and the inability of users to verify the provenance of AI-generated claims. To resolve this, we developed an integrated platform that prioritizes mechanistic interpretability through two primary channels: (i) structural evidence grounding and (ii) interactive topological visualization. Unlike conventional chat interfaces that provide monolithic responses, our platform decomposes the reasoning process into a verifiable audit trail, allowing clinical researchers and oncologists to scrutinize the logic underlying every therapeutic or mechanistic assertion.

A central feature of the interface is the **Evidence Provenance Module**. Every synthesized response is intrinsically linked to the 100K-article full-text hypergraph, with specific assertions mapped to their originating sources via persistent identifiers (e.g., [PMC7352562], [PMC7358280]). This granular level of citation ensures that the “Mechanistic Reasoning” identified as a barrier in current medical LLMs is transformed into a navigable bibliography. By selecting a specific segment of the generated answer, users can instantly access the supporting literature segments, thereby facilitating rapid manual verification during time-sensitive scenarios such as Tumor Board reviews.

The platform’s **Dynamic Evidence Graph** provides a real-time topological representation of the reasoning path. As the LLM synthesizes an answer, the system simultaneously renders an interactive vertical flow of the involved hyperedges and *n*-ary relationships. This visualization moves beyond simple node, link diagrams by representing complex biological assemblies, such as the Tryptophan-Kynurenine-AhR axis, as structural entities. This allows oncologists to visually confirm that the proposed mechanism respects known biological hierarchies and does not rely on stochastic associations.

## 3 Discussion

The development of **MetaKnogic-Alpha** represents a fundamental shift from probabilistic text synthesis to structured, mechanistic reasoning in biomedical AI. By bridging the gap between unstructured, large-scale literature and deterministic biochemical logic, our system addresses the critical “probabilistic bottleneck” that has historically limited the clinical utility of general-purpose foundation models. While models like GPT-5.1 exhibit remarkable fluency, their lack of structural grounding frequently results in biochemically implausible assertions. MetaKnogic-Alpha resolves this by enforcing a hyper-relational evidence audit trail, achieving high accuracy in scenarios where direct literature evidence is available.

The impact of this work on biomedical knowledge discovery is two-fold. First, it demonstrates that the “mechanistic dark matter” hidden within the results sections of full-text articles can be systematically extracted and organized into a million-scale hypergraph at a fraction of the traditional cost. Our optimized pipeline provides a scalable blueprint for small-to-medium-sized research laboratories to build expert-level knowledge bases without the need for prohibitive industrial-grade compute clusters. Second, by integrating a structured metabolic network as a logical constraint layer, we have shown that LLMs can be “steered” to respect biological first principles, such as mass balance and reaction directionality, thereby providing a safeguard against stochastic hallucinations in high-stakes clinical contexts.

Despite its performance, several limitations of the current architecture warrant consideration. While the hypergraph provides a rigorous evidence base, the final synthesis still relies on the generative capabilities of external reasoning engine (e.g., GPT-4/5.1). This dependency introduces a potential “black-box” element in the final narrative construction, although our interactive visualization of evidence paths significantly mitigates this by allowing manual verification of every hyper-relation.

Looking forward, the computational extensibility of MetaKnogic-Alpha allows for a clear path toward “million-scale” knowledge synthesis. We envision extending the current foundation to encompass the entirety of the PubMed Central open-access repository, creating a universal mechanistic engine for all known biological interactions. Beyond literature, the next frontier for MetaKnogic-Alpha lies in the integration of spatial transcriptomics and complex single-cell multi-omics data. Incorporating the spatial architecture of the tumor microenvironment into the hypergraph would allow for the modeling of paracrine signaling and metabolic competition at a resolution previously unattainable by text-only or bulk-data systems.

Finally, the MetaKnogic-Alpha framework provides a foundational infrastructure for the “Digital Twin” approach in precision medicine. By constructing integrative, patient-centered knowledge graphs that synthesize a patient’s clinical records with the global breadth of scientific literature, the system enables the generation of case-specific agents. These agents do not merely summarize data; they act as mechanistic reasoning partners for Tumor Boards, identifying personalized therapeutic vulnerabilities grounded in verifiable biological logic. As we continue to refine this architecture, we anticipate that MetaKnogic-Alpha will serve as a cornerstone for the transition from descriptive to predictive artificial intelligence in clinical oncology.

## 4 Methods

### 4.1 Strategic Corpus Prioritization and Mechanistic Density Optimization

To establish a high-fidelity knowledge substrate for **MetaKnogic-Alpha**, we developed a multi-factorial prioritization protocol designed to maximize mechanistic density while addressing the inherent informational redundancy in biomedical literature. Starting from an initial pool of over 7 million PMC full-text articles, we observed that while the volume of metabolic literature is vast, the core biochemical logic is often recapitulated across numerous redundant reports[15, 37, 38].

To ensure the most representative and reliable evidence was prioritized for the “alpha” release, we applied a curated dictionary of 3,400 domain-specific terms and implemented a composite importance scoring system which integrates text-evidence strength, citation impact, recency, and journal prestige. This allowed us to distill the corpus into a 100K-paper “discovery backbone” that maintains broad coverage across the entire metabolic landscape without the computational noise of redundant or low-signal publications.

Critical to our approach, we monitored keyword coverage across importance percentiles to verify that our prioritization did not result in “topic collapse,” ensuring that even rare regulatory nodes and distal metabolic links were preserved. We frame this primary release as a scalable architectural blueprint, prioritizing the quality of the “Knogic” (Knowledge-Logic) over mere document volume. A comprehensive description of the scoring algorithms, ambiguous term controls, and percentile-based coverage validation is provided in the **Appendix A**.

### 4.2 Large-Scale Full-Text Literature Ingestion and Structural Chunking

The knowledge base of MetaKnogic-Alpha is founded upon the ingestion of over 100K full-text articles retrieved from the PMC. To resolve the extreme computational latency inherent in processing million-scale full-text corpora, we designed a high-concurrency, asynchronous batch processing pipeline. This architecture facilitates a several-hundredfold increase in ingestion throughput, compressing a projected multi-month ingestion timeline into approximately hours, while simultaneously reducing API operational expenditures by 50%.

Documents are subjected to a “Structural Semantic Chunking” protocol that avoids the information loss associated with fixed-length windowing. Utilizing a rule-based parser, the system identifies natural semantic boundaries, such as section headers and paragraph terminations, to ensure that high-order biological narratives remain contextually intact[39, 40]. This pre-processing layer is vital for delivering high-density, semantically complete data segments to the downstream extraction engine.

### 4.3 Automated N-ary Hyperedge Instantiation from Full-Text Literature

While the theoretical advantages of hypergraphs for representing complex biological systems have been established[41, 42], the automated construction of such structures from unstructured text remains an open challenge[27, 43]. A primary contribution of the MetaKnogic-Alpha pipeline is the high-throughput transition from traditional binary edge extraction to n-ary hyperedge instantiation across a million-scale document corpus.

Unlike standard knowledge graph extraction protocols that decompose biochemical reactions into independent, pairwise entity associations (e.g., Enzyme-Substrate, Enzyme-Product), our agentic framework is designed to capture the intrinsic multi-entity nature of biological events. By identifying overlapping semantic clusters within full-text segments, our extraction agents group all participating substrates, products, and co-factors into single, unified hyperedges. This method preserves the stoichiometric and topological integrity of the evidence, which we demonstrate is essential for resolving deep mechanistic questions that simple binary graphs, and standalone LLMs like GPT-5.1 fail to address.

#### 4.3.1 Biochemical Grounding via Metabolic Network Integration

To enforce biochemical consistency during retrieval and synthesis, we integrated a curated metabolic multigraph comprising several thousand reactions and compounds, represented as typed relations among **genes, reactions**, and **metabolites**[3]. This deterministic framework is indexed for efficient access and serves as a logical grounding layer that anchors the generative process in mechanistic structure. Reaction nodes carry curated attributes such as pathway membership, compartment, directionality (reversible/irreversible), and reference identifiers (e.g., KEGG reaction IDs), while metabolite nodes encode compound identifiers and compartment-specific forms.

Every literature-extracted entity is subjected to a Structural Anchor strategy that maps mentions to canonical nodes in the metabolic graph (including normalization and alias handling). Once anchored, candidate relationships are cross-verified against allowable graph semantics, requiring support by explicit edges (e.g., gene→reaction catalysis; metabolite→reaction consumption; reaction→metabolite production) and respecting reaction directionality and compartment constraints. At runtime, the anchored seeds enable deterministic neighborhood retrieval and subgraph construction (followed by local “reaction-closure” to include co-substrates, products, and catalyzing genes), which reduces unsupported mechanistic claims and improves traceability of generated explanations to biochemically valid connections.

### 4.4 Hierarchical Knowledge Discovery and Mechanistic Synthesis

The retrieval and synthesis architecture of **MetaKnogic-Alpha** is designed to mirror the multi-layered reasoning process required for complex metabolic research[44, 45, 46, 47]. Rather than utilizing a single-shot retrieval approach, the system employs a three-tiered “Hierarchical Discovery” protocol: agentic query enrichment, multi-hop topological expansion, and deterministic metabolic grounding. This ensures that the final synthesis is not only semantically relevant but also biochemically consistent with the known laws of cellular metabolism.

#### 4.4.1 Tier 1: Agentic Query Enrichment and Semantic Discovery

To bridge the gap between abstract user inquiries and the high-resolution nomenclature of the million-scale hypergraph, the process initiates with an “Agentic Enrichment” phase. A specialized biomedical agent parses the raw query to identify latent biological entities, enzymatic participants, and pathway involvement[48, 49, 50]. This enriched query is then projected into a 768-dimensional latent space via the embedding model. By identifying dense semantic “seeds” within the vector store, the system ensures that the entry points into the knowledge base are contextually precise, effectively filtering out lexical noise that frequently plagues keyword-based retrieval[51].

#### 4.4.2 Tier 2: Multi-Hop Topological Expansion

The core structural strength of MetaKnogic-Alpha is the transition from textual retrieval to “Topological Discovery”. For each identified semantic seed, the system performs an automated multi-hop breadth-first traversal within the hypergraph store.

This expansion retrieves first- and second-order neighbors, including shared metabolites and regulatory co-factors that may be disjointed across the 100K-article corpus. This step is essential for resolving non-linear pathways, as it provides the generative model with a “global mechanistic neighborhood” rather than isolated text fragments[28, 27].

#### 4.4.3 Tier 3: Deterministic Metabolic Grounding and Synthesis

To mitigate the risk of probabilistic hallucinations[52, 53], all retrieved hyper-relations are subjected to a “Biochemical Sanity Check” against a deterministic metabolic network. The system cross-references literature-derived links against established metabolic reaction templates. This hierarchical filtering ensures that the output is both evidence-based and biologically plausible, providing a verifiable audit trail from the resulting assertion back to the original literature.

##### Algorithm 1 Hierarchical Discovery and Mechanistic Synthesis

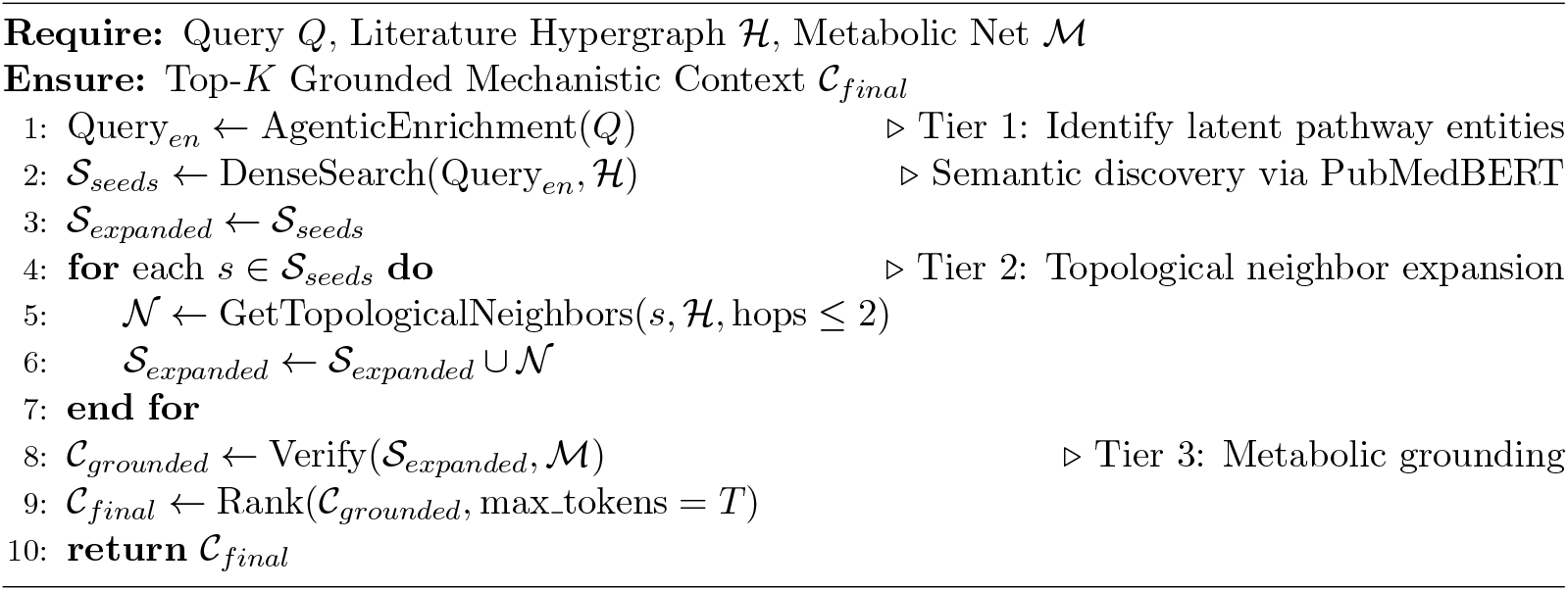

## 5 Implementation and Code Availability

The framework is implemented as a high-performance, distributed service architecture designed for million-scale knowledge synthesis. The system is written in Python and utilizes an asynchronous, non-blocking framework to achieve a several-hundredfold reduction in ingestion latency, effectively compressing a projected multi-month processing timeline into several hours.

### 5.1 Architectural Framework and Computational Stack

The core logic and neural processing modules are built using PyTorch, facilitating the high-throughput generation of 768-dimensional embeddings via the ModernPubmedBERT model. The system utilizes a hybrid, multi-modal database architecture to manage the complexity of biological hypergraphs and metabolic pathways:

- **Metabolic Network and Topological Analysis:** We utilized **NetworkX** to construct and maintain the deterministic metabolic grounding layer. This in-memory graph representation allows for sub-millisecond path-finding and stoichiometric validation, ensuring that all literature-extracted hyper-relations are consistent with established biochemical laws before being synthesized into a final response.
- **Hypergraph Storage and Relational Logic: Neo4j** serves as the primary persistent storage for the million-scale hypergraph. It enables complex, multi-hop Cypher-based traversals across the 100K-article corpus, providing the structural backbone for the system’s topological neighbor expansion.
- **High-Dimensional Vector Retrieval:** A **Milvus** cluster provides the vector similarity search layer, enabling high-speed retrieval of dense semantic candidates based on the 768-dimensional embeddings.
- **Serving Infrastructure:** The user interface and API layer are powered by **FastAPI** and **Uvicorn**, providing a high-concurrency gateway for real-time mechanistic querying and evidence visualization.

The pipeline is containerized and deployed on a high-performance computing (HPC) cluster equipped with NVIDIA A100 GPUs. This hardware acceleration, combined with the asynchronous batch processing engine, facilitates a 50% reduction in operational costs by maximizing hardware utilization and minimizing API overhead.

### 5.2 Code Availability and Open Access

In accordance with open-science principles, the full computational suite and the resulting knowledge substrate are publicly available under the MIT License.

- **Core Repository:** The complete source code is available at: https://github.com/paveethrans/MetaKnogic-Alpha.
- **Interactive Discovery Portal:** An interactive web application for exploring the evidence-based audit trails and metabolic mappings can be accessed at the project website.

## A Strategic Corpus Prioritization and Mechanistic Density Optimization

### A.1 Relevance-based corpus prioritization for full-text ingestion

As a prerequisite for scalable full-text hypergraph construction, we first established a ranked candidate literature pool using the composite relevance score described in Methods (Eq. 4). The relevance-score statistics in Fig. 2 and Fig. 3 are computed on the 1M+ candidate corpus used for backbone selection. Fig. 2 shows the retained fraction as a function of the threshold, providing a controllable trade-off between corpus size and expected relevance. In parallel, Fig. 3 verifies that high-relevance strata preserve broad keyword coverage rather than collapsing onto a small set of highly frequent terms. Using this ranked pool, we selected an initial operating point for end-to-end extraction, yielding the 100K full-text subset used for hyperedge instantiation and subsequent system benchmarking.

**Figure 2.**
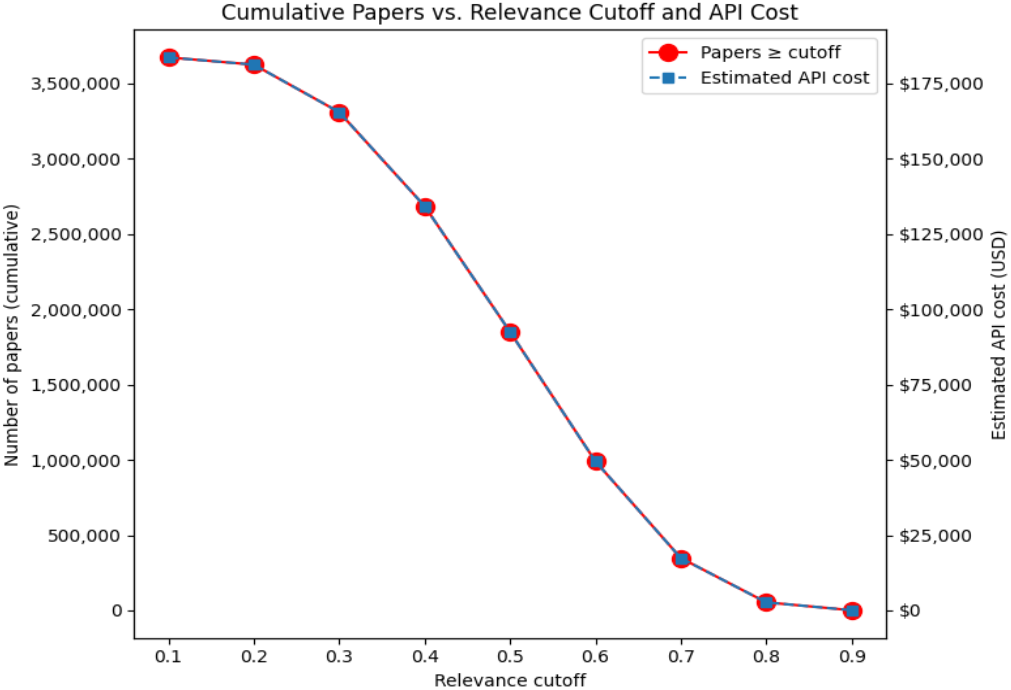
Fraction of articles retained as a function of the relevance-score threshold (1M+ corpus)

**Figure 3.**
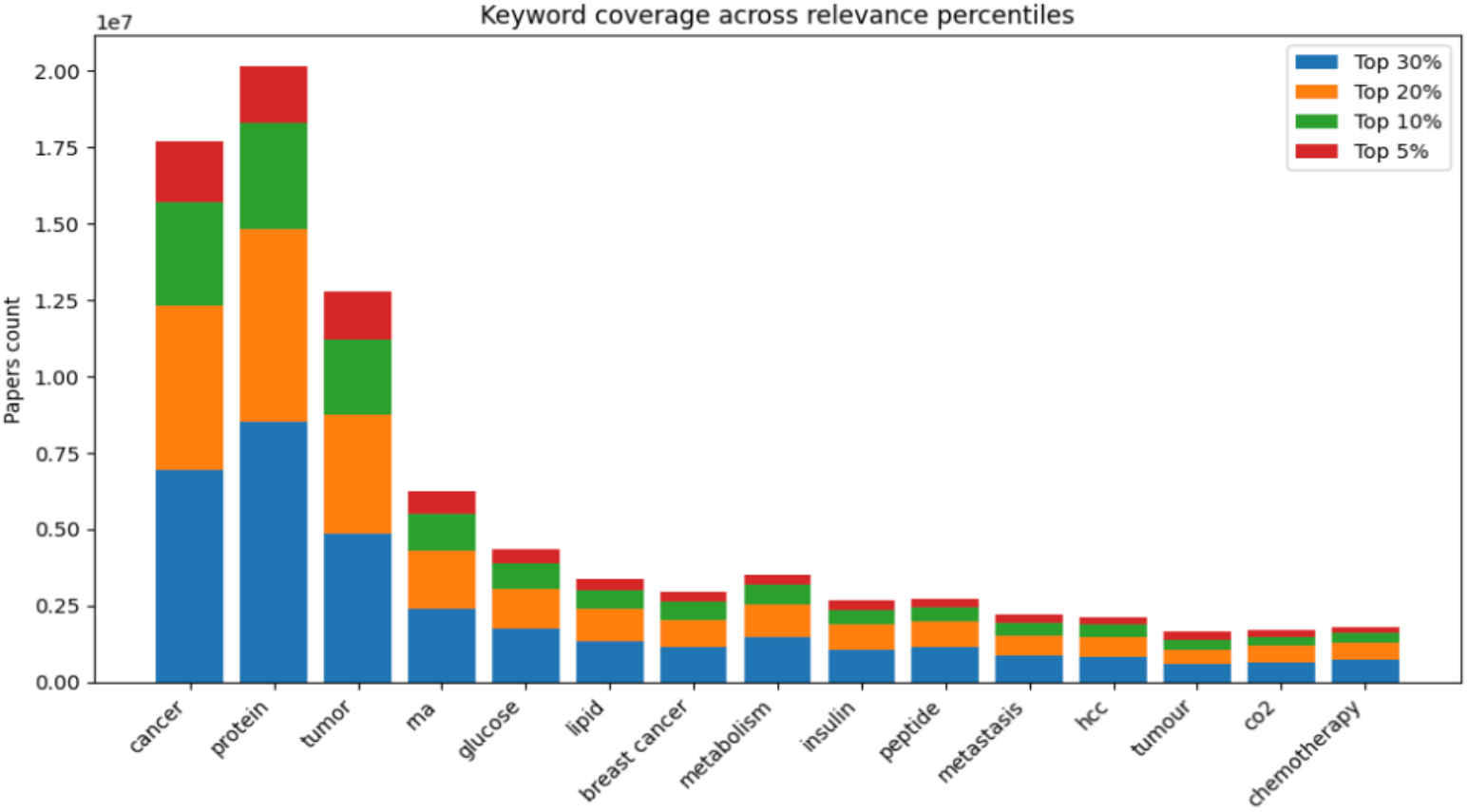
Keyword coverage across relevance-score percentiles

### A.2 Literature ingestion and paper selection

To construct an evidence-grounded corpus for hypergraph extraction at million-scale, we paired PMC full-text with OpenAlex metadata and applied term-based filtering followed by relevance-score ranking.

#### A.2.1 Paper ingestion, alignment, and term-based filtering

We ingested approximately 7 million PMC full-text articles (excluding author manuscripts) and linked each record to OpenAlex work-level metadata (e.g., cited-by counts, publication year, and venue identifiers). Records were aligned primarily by PMCID (with secondary cross-identifiers where available), yielding 6.6 million matched OpenAlex–PMC pairs used for downstream filtering and ranking.

To focus on mechanistically relevant biomedical literature, we applied a curated dictionary of 3,400 domain keywords and synonyms using exact matching with word-boundary constraints and controlled normalization. Papers containing at least one dictionary term were retained, yielding 5.5 million candidate articles.

#### A.2.2 Relevance score design and filtering

The 5.5 million term-matched papers remained too large for exhaustive full-text extraction. We therefore computed a composite relevance score combining (i) text evidence strength, (ii) citation impact, (iii) recency, and (iv) journal prestige. Each signal was mapped to a robust [0, 1] scale via quantile ranking, denoted qrank01(·) to reduce sensitivity to outliers.

##### Text signal

The purpose of this signal is to quantify strength of an article based on the positions of keywords present; We compute two text-based components: a section-weighted match score *T*_1_ and a keyword-density score per 1,000 tokens *T*_2_.

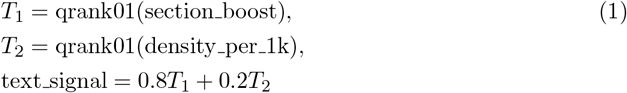

Where section boost upweights dictionary matches found in high-salience sections (e.g., title, abstract, results) relative to low-salience sections (e.g., methods, references). And density per 1k is the number of dictionary matches per 1,000 tokens and was upper-bounded to suppress documents dominated by repetitive matches.

##### Citation signal

The purpose of this signal is to act as an impact proxy and quantify the reliability of an article’s content. Where `cited_by_count` stores the number of times the article was cited and referred.

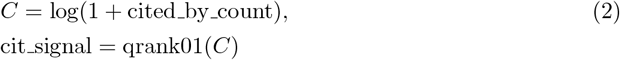

##### Journal prestige signal and recency signal

The purpose of this signal is to mildly prioritize recent articles by a small margin alongside the journal’s prestige score (see Algorithm 2), where `pub_year` points to the publication year, base year is set as xxxx.

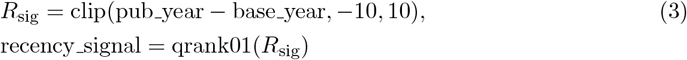

###### Algorithm 2 Journal prestige signal computation

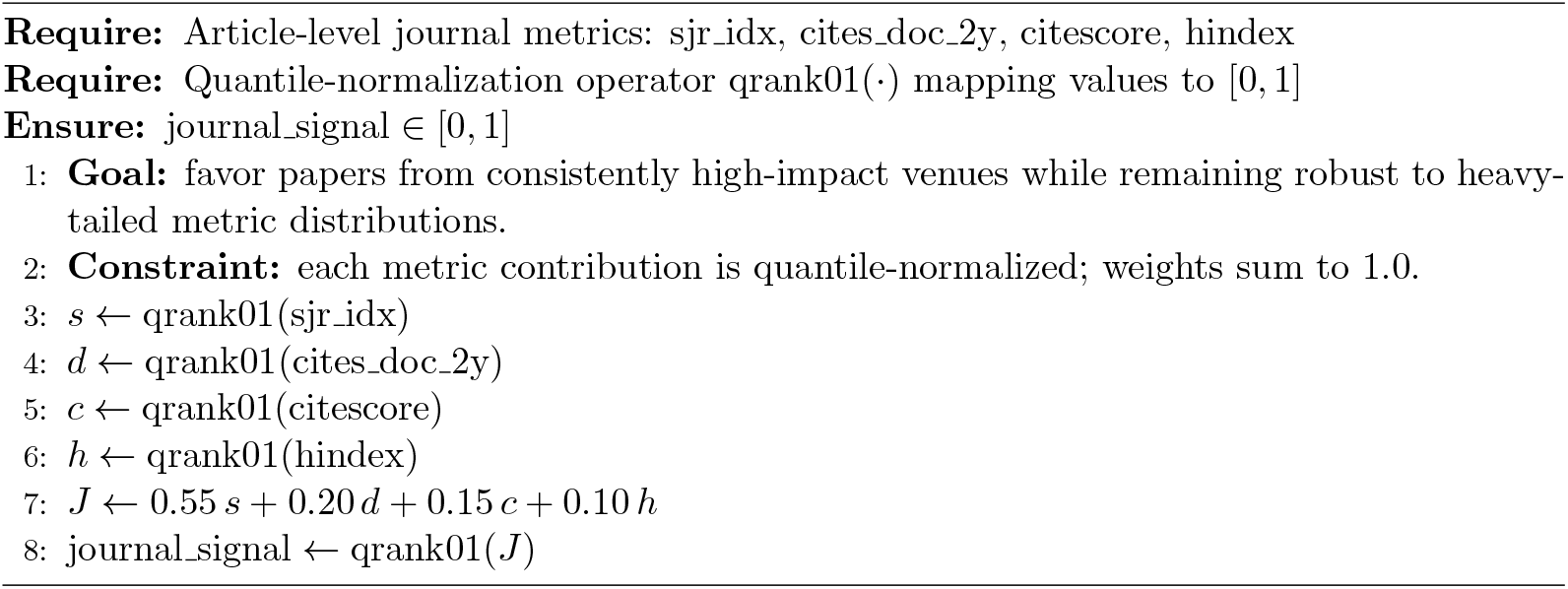

##### Total relevance score

Together with all previously computed signal values, the total relevance score was used to filter low-priority papers and to rank the remaining corpus for computationally expensive full-text extraction and subsequent graph and hyperedge construction.

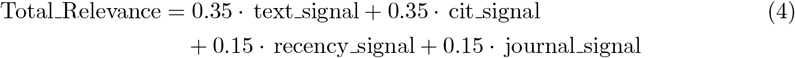

#### A.2.3 Ambiguous synonym control and case-sensitive safeguards

Short synonyms (3–4 characters) can inflate matches and degrade precision. For example, ALL (acute lymphoblastic leukemia) produced *>* 5 million matches when applied naively; removing this token reduced the candidate set to *<* 3.8 million while preserving the long-form concept (approximately 13,000 papers).

We mitigated such collisions using a curated stoplist of ambiguous short synonyms, strict word-boundary constraints for high-risk tokens, and case-sensitive matching to suppress false positives from uppercase table artifacts and boilerplate.

#### A.2.4 Resulting backbone corpus

After term filtering, relevance-score ranking, and ambiguity controls, we obtained a 3.5 million paper backbone corpus used for downstream knowledge hypergraph construction. In selecting the backbone, we prioritized breadth of keyword coverage across the 3,400-term dictionary (including gene-name coverage) rather than depth on a small set of highly frequent terms.

We sorted papers by Total Relevance (Eq. 4) and monitored keyword coverage across relevance-score percentiles to ensure that high-ranked strata did not collapse onto a narrow subset of terms (Fig. 3). We used the relevance-score cutoff curve to choose tractable thresholds for computationally expensive full-text extraction (Fig. 2). From the 3.5M ranked papers, an initial threshold of Total_Relevance *>* 0.7 yielded an approximately 100K-paper corpus used for end-to-end extraction and evaluation, with planned expansion toward 1M+ papers as computational resources permit.

